# Revealing Chloroquine’s Antimalarial Mechanism: The Suppression of Nucleation Events during Heme to Hemozoin Transformation

**DOI:** 10.1101/2023.06.15.545078

**Authors:** Rahul Singh, Rashmi Singh, Velaga Srihari, Ravindra D. Makde

**Author notes:** Corresponding Author: Dr. Ravindra D. Makde, Beamline Development, and Application Section, Bhabha Atomic Research Centre, Mumbai 400085. India.

## Abstract

Malaria parasites generate toxic heme during hemoglobin digestion, which is neutralized by crystallizing into inert hemozoin (β-hematin). Chloroquine blocks this detoxification process, resulting in heme-mediated toxicity in malaria parasites. However, the exact mechanism of chloroquine’s action remains unknown. This study investigates the impact of chloroquine on the transformation of heme into β-hematin. The results show that chloroquine does not completely halt the transformation process but rather slows it down. Additionally, chloroquine complexation with free heme does not affect substrate availability or inhibit β-hematin formation. SEM and XRD studies indicate that the size of β-hematin crystal particles and crystallite increases in the presence of chloroquine, suggesting that chloroquine does not impede crystal growth. These findings suggest that chloroquine delays hemozoin production by perturbing the nucleation events of crystals and/or the stability of crystal nuclei. Thus, contrary to prevailing beliefs, this study provides a new perspective on the working mechanism of chloroquine.

## INTRODUCTION

Despite the recent progress in the fight against malaria, this disease is still responsible for annually 247 million cases in 82 malaria endemic countries [1]. Furthermore, the efficacy of frontline antimalarial drugs is already under question due to widespread chloroquine (CQ) resistance [2] and the emergence of artemisinin resistance [3]. The development of future antimalarial under such circumstances demands investigating the critical physiological processes of *Plasmodium*. The clinical symptoms of malaria are due to the asexual multiplication of the *Plasmodium* inside the host RBC [4]. During this stage, the *Plasmodium* digests about 75% of hemoglobin inside the infected erythrocyte [5]. Within the parasite, this massive hemoglobin digestion happens within a subcellular compartment of the digestive vacuole, liberating copious amounts of free heme [6]. Free heme is considered highly cytotoxic [7]. Without the active heme oxygenase gene [8], *Plasmodium* detoxifies 90-95% of the host-derived heme by converting it into an inert hemozoin crystal (synthetic form is β-hematin) [9] [10]. Hence, hemozoin (Hz) production is the only way to sequester free heme and is therefore considered a physiological bottleneck in malaria parasites [11]. Several theories have been proposed in *Plasmodium* to explain the large-scale heme to Hz conversion. Generally, these theories either support spontaneous transformation [12]-[13], lipid-mediated [14][15][16], or protein-mediated conversion of heme into Hz [17][18][19][20][21]. Among all known mediators of Hz production, heme detoxification protein (HDP; UniProt ID: Q8IL04) is shown to be most efficient under in-vitro settings and proven indispensable for the parasite’s viability [19].

Antimalarials derived from quinolone were the first line of drugs used to treat malaria. Even after the development of drug resistance, CQ is still used with artemisinin and other medicines to treat malaria. There are several hypotheses about the working mechanism of CQ [22]. However, CQ is primarily reported to interfere with the Hz production process in *Plasmodium* [23][24][25][26][27][28][29][30] for inducing toxicity to the parasite. There are two primary theories about how CQ perturbs Hz production in *Plasmodium*. As per the first opinion, the quinolone family of drugs forms a complex with free heme making it unavailable for Hz formation [23][24][25][26]. The recent second opinion is about inhibiting Hz crystal growth via binding CQ on the crystal surface and inhibiting further substrate addition [27][28][29][30]. Even after decades of work, a consensus about the CQ mode of action has yet to be achieved. The sequestration of heme into Hz is a realistic target for developing new antimalarials. Therefore, the mechanism of inhibiting Hz production by antimalarials like chloroquine may facilitate future drug development.

In this in-vitro study, by using HDP as a catalyst, we tried to imitate the native *in-vivo* conditions during heme to Hz transformation. The influence of CQ on the conversion of heme into β-hematin was studied by inhibition, time-kinetics, XRD, and SEM experiments. As a result, our study evaluates the existing beliefs on the working mechanism of CQ and presents a new aspect of this subject.

## RESULT

### CQ-mediated inhibition

To study the CQ’s role in inhibiting heme to β-hematin transformation, we performed the inhibition studies at pH 4.8 and 5.2. The experiments were performed in series with the same reaction condition, except for different heme substrate concentrations. The reaction at pH 5.2 was stopped arbitrarily at ∼50% completion of the reaction (Fig. S1A). However, all reaction series at pH 4.8 were stopped at two discrete time points (Fig 1A), one well before the completion of the reaction (5 min.; Fig. 1B) and the other just close to the completion of the reaction (7 min. Fig. 1C). These specific time points were selected to ensure that the reaction dose not reach completion so that we could make observations during the active phase of heme to Hz transformation (Fig 1A). Both experiments at different pH values yielded distinct IC_50_ values of CQ for reaction series with different heme concentrations (Fig. 1B & C; Table S1, Fig S1A; Table S2). We observed different values of CQ IC50 for different heme concentrations and reaction times. The range of CQ IC_50_ value is between 11-46 μM (Fig. 1D). We observed a positive trend between the IC50 value and the reaction’s heme substrate concentration (Fig 1D and S1B). We also observed that the efficiency of CQ to inhibit β-hematin inhibition is better at 5 min than at 7 min. Therefore there is a positive trend between the IC50 value and the reaction time (Fig.1D).

**Figure 1:**
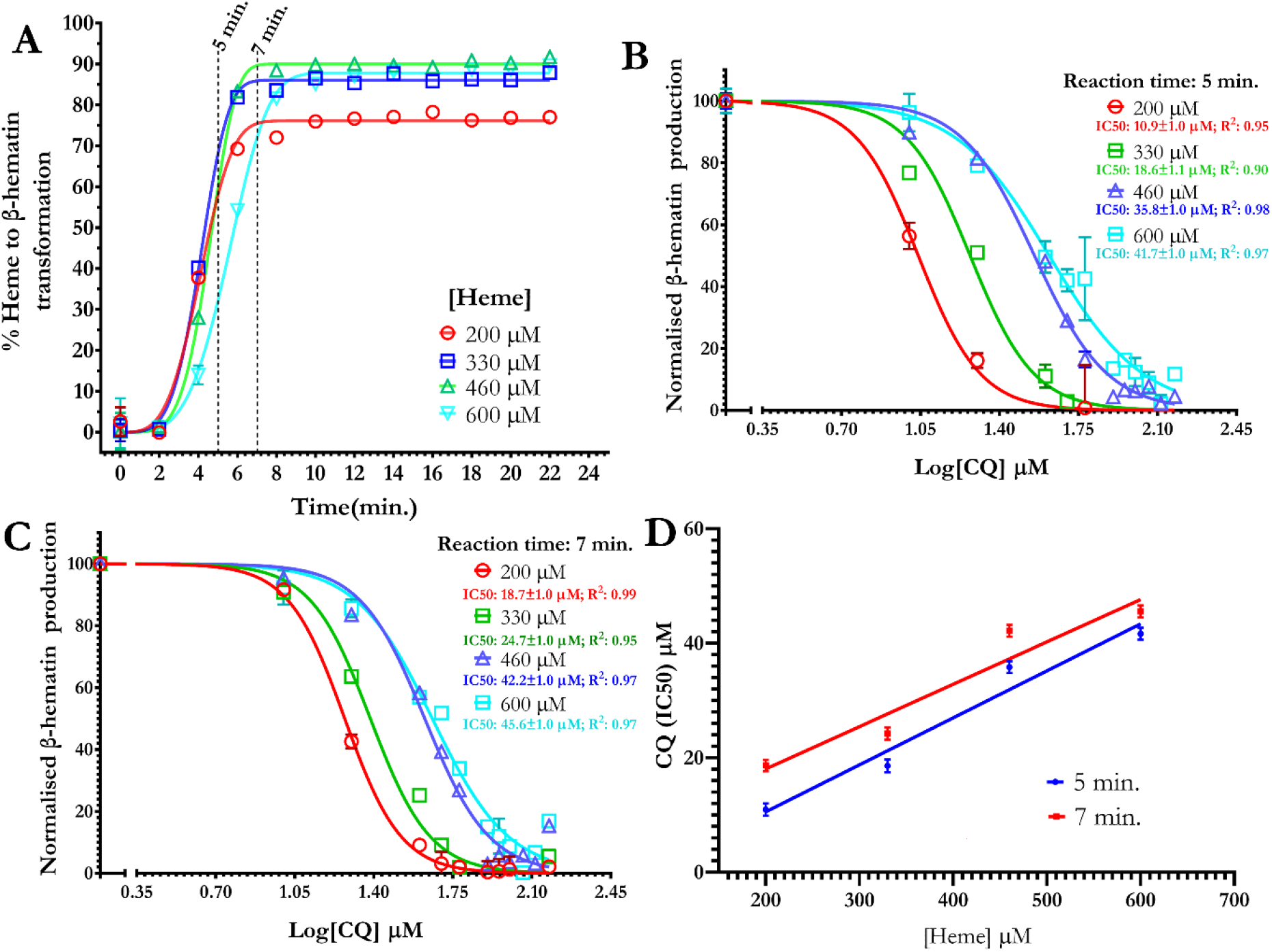
CQ mediated inhibition of β-hematin production (pH4.8). *A*, Change in normalized β-hematin production as a function of time in reaction series with the constant HDP (1.5 μM) and color-coded heme concentration. The dotted line represents the time point of 5 and 7 min. *B&C*, The graph represents a change in normalized β-hematin production as a function of CQ concentration at the constant HDP and color-coded heme concentrations. Individual points (square, circle, or triangle) represent the mean value calculated by measuring the change in free-heme concentration present in the assay supernatant. The bar represents the error between replicate reactions (N=2). The fitting statistics is given in Table S1. *D*, The graph shows a positive trend between the reaction’s CQ IC50 value and heme concentration.

### Time-kinetics

We performed time-kinetics experiments in reaction series with the same reaction composition, except for a single constituent with varying concentrations among series. The amount of β-hematin (product) produced in a reaction was assessed by measuring the amount of spent heme (substrate) via pyridine hemochromogen assay. The role of HDP (Fig. 2A, S2A) and CQ (Fig. 2B, S2B) on heme to β-hematin transformation were analyzed with this method. In addition, the effect of introducing CQ at different time points during time kinetics was also analyzed to capture the precise role of CQ in inhibiting β-hematin production (Fig. 2C, S2C). All time-kinetic experiments were performed at pH 4.8 (Fig. 2) and 5.2 (Fig. S2), during which the percentage of heme transformed to β-hematin was monitored in reference to the reaction time. The time-kinetic data were analyzed by Kolmogorov-Johnson-Mehl-Avrami (KJMA) equation [31][32]. Sometimes the fitting of our time-kinetic data into the KJMA equation yielded a value of the Avrami constant (n) of more than 4 (Table S3 & S4). This observation is unusual as “n” generally takes an integer value between 1 and 4 [33]. Avrami constant defines the nature of crystal growth, and as earlier observed [34], sometimes its value exceeds 4. In our case, we believe this is due to the heterogeneous nature of the reaction mixture due to heme aggregates. Under suitable conditions, these aggregates could influence or become a site for crystal nucleation and growth, adding more variables in phase transformation and causing the n value to exceed 4. Since, Avrami constant has not been used in any way during the analysis and the interpretation of the data, we have not discussed this anomaly further in the manuscript.

**Figure 2.**
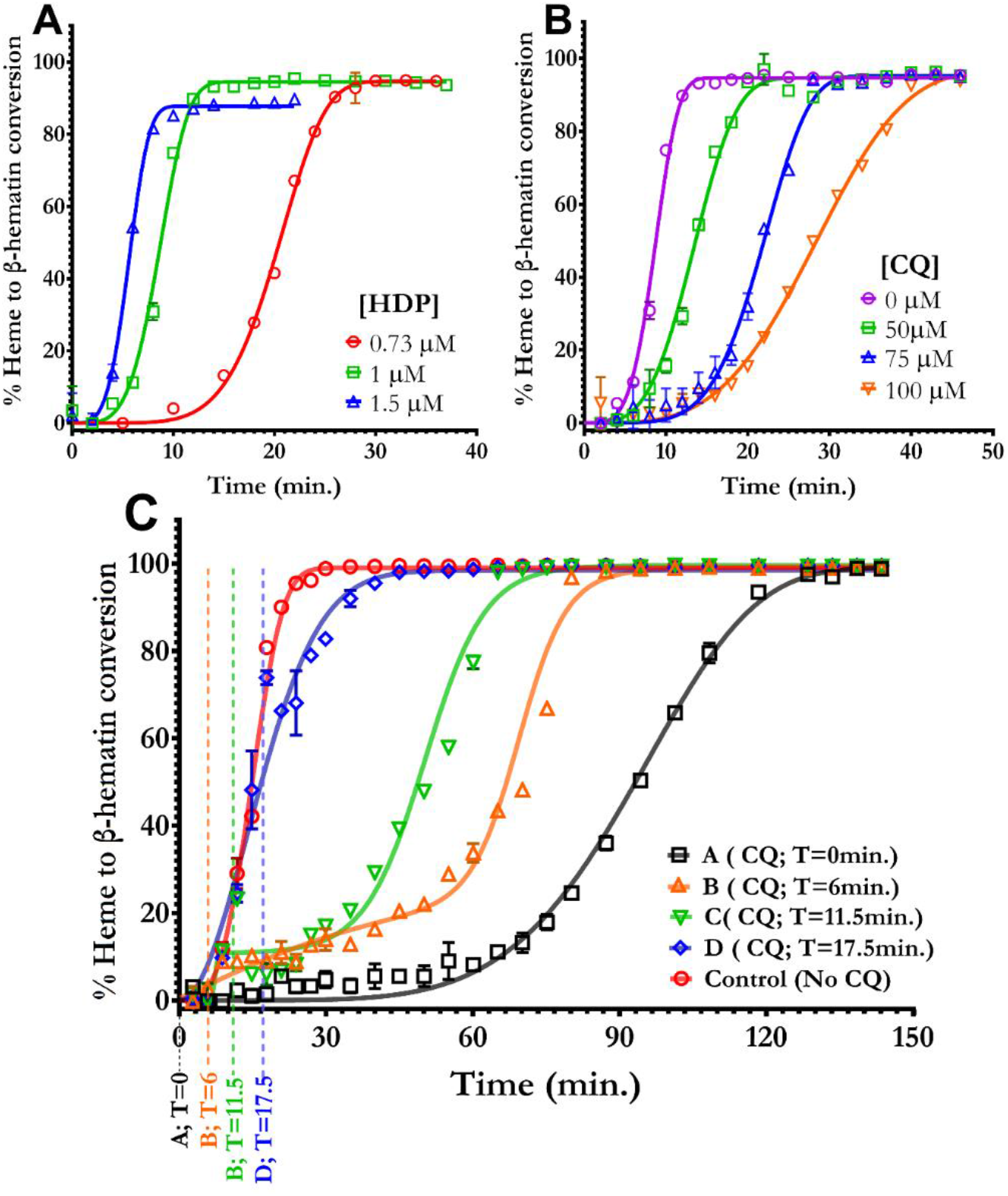
Time-kinetics at pH 4.8. ***A***, The graph shows the change in the percentage of heme transformed to β-hematin as a function of time at constant substrate (600 μM) concentration and color-coded series with different HDP concentrations. ***B***, Percent heme to β-hematin transformation as a function of time at a constant concentration of HDP (1 μM) and heme substrate (600 μM), but different CQ concentrations (color-coded) in each reaction series. ***C***, The graphs represent a change in normalized β-hematin production as a function of time at the constant HDP (0.8 μM) and heme (600 μM) concentrations. Each reaction except the control series was exposed to 100 μM CQ at a different time point after the onset of the reaction, i.e., 0 min. (series A), 6 min. (series B), 11.5 min. (series C) and 17.5 min. (series D) represented by color-coded dotted lines and arrows. The line represents the best fit to the KJMA equation or biphasic sigmoid growth equation (color-coded). Individual points (square, circle, or triangle) represent the mean value calculated by measuring the change in free-heme concentration present in the assay supernatant. The bar represents the error between replicate reactions (N=2). The fitting statistics are given in Table S3.

In all time-kinetic experiments, the heme to β-hematin transformation followed a sigmoid path with reaction time, characteristic of phase transformation reaction. Depending on the variable constituent’s concentration, significant differences in the relative profile of the sigmoid curve among reaction series were observed (Fig. 2 & S2). In reaction series with varying HDP concentrations (Fig 2A & S2A), the initial asymptotic stage decreases with increasing HDP concentration. In addition, the reaction completion time increases with a decrease in HDP concentration (Fig. 2A & S2A). At the stationary stage, the percent conversion of heme substrate into the β-hematin product also slightly varies for different HDP concentrations.

Like HDP, the role of CQ is also studied via the time-kinetic experiment in reaction series with the same concentration of heme and HDP but different CQ concentrations (Fig. 2B & S2B). We observed that all reaction series reached completion (∼95% substrate depletion) irrespective of the presence of CQ in the reaction mixture. However, opposite to the effect of HDP, the reaction series with a higher CQ concentration required more time to achieve completion. Similarly, contrary to HDP’s effect, the initial asymptotic stages gradually become more pronounced, with increasing CQ concentration (Fig. 2B & S2B). Under low concentration of HDP, the ability of CQ to increase the duration of an initial asymptomatic stage and overall time of transformation reaction is particularly pronounced (Control v/s series A; Fig. 2C & S2C). The observed decrease in completion time with increasing HDP concentration suggests its active involvement in catalyzing the transformation reaction. In contrast, the increase in completion time with increasing CQ concentration indicates the slowing down of the reaction. Additionally, we have highlighted that the endpoint of the transformation reaction remained unchanged irrespective of the presence of CQ in the reaction mixture.

Sometimes, the introduction of CQ after the onset of the transformation reaction can induce bi-phasic growth (Fig. 2C, series B&C). Therefore, the influence of CQ on the duration of the initial asymptomatic stage might be difficult to quantify. However, the quantification of reaction completion time is relatively easy. The effect of CQ on reaction time and the duration of the initial asymptomatic stage depend on the CQ introduction time point after the onset of the reaction. That is., if CQ is introduced into the reaction mixture at the beginning of the reaction (Fig 2C, Series A), it takes ∼110 minutes more than the control reaction to complete. This delay in the reaction completion can be decreased by ∼50% (110 to 57 min.) by just a mere delay of 6 minutes (Fig 2C, Series B) to introduce CQ into the reaction mixture after the onset of the reaction. Therefore, the ability of CQ to influence heme to β-hematin transformation decreases with the delay in introducing CQ into the reaction mixture after the onset of the reaction (Fig. 2C & S2C).

### CQ pull-down assay

In the present study, we employed a CQ pull-down assay to investigate the interaction between CQ and heme during the heme to β-hematin transformation. The acidic reaction environment of heme to β-hematin transformation causes the heme substrate to become insoluble, while CQ is soluble. However, upon complex formation between CQ and heme, the CQ in complex with heme also becomes insoluble and can be precipitated along with heme via centrifugation. To quantify the amount of uncompleted CQ left in the reaction supernatant, we measured the absorbance at 329 nm (extinction coefficient of 16.6 mM^-1^.cm^-1^). The CQ precipitation capacity for 600 μM heme was found to increase with the mixing time (Fig S3A), and we allowed the heme substrate and CQ to interact with each other for 1 hour via vigorous shaking before initiating the transformation reaction by adding HDP (1μM) to the reaction mixture (Fig. 3A)

**Figure 3:**
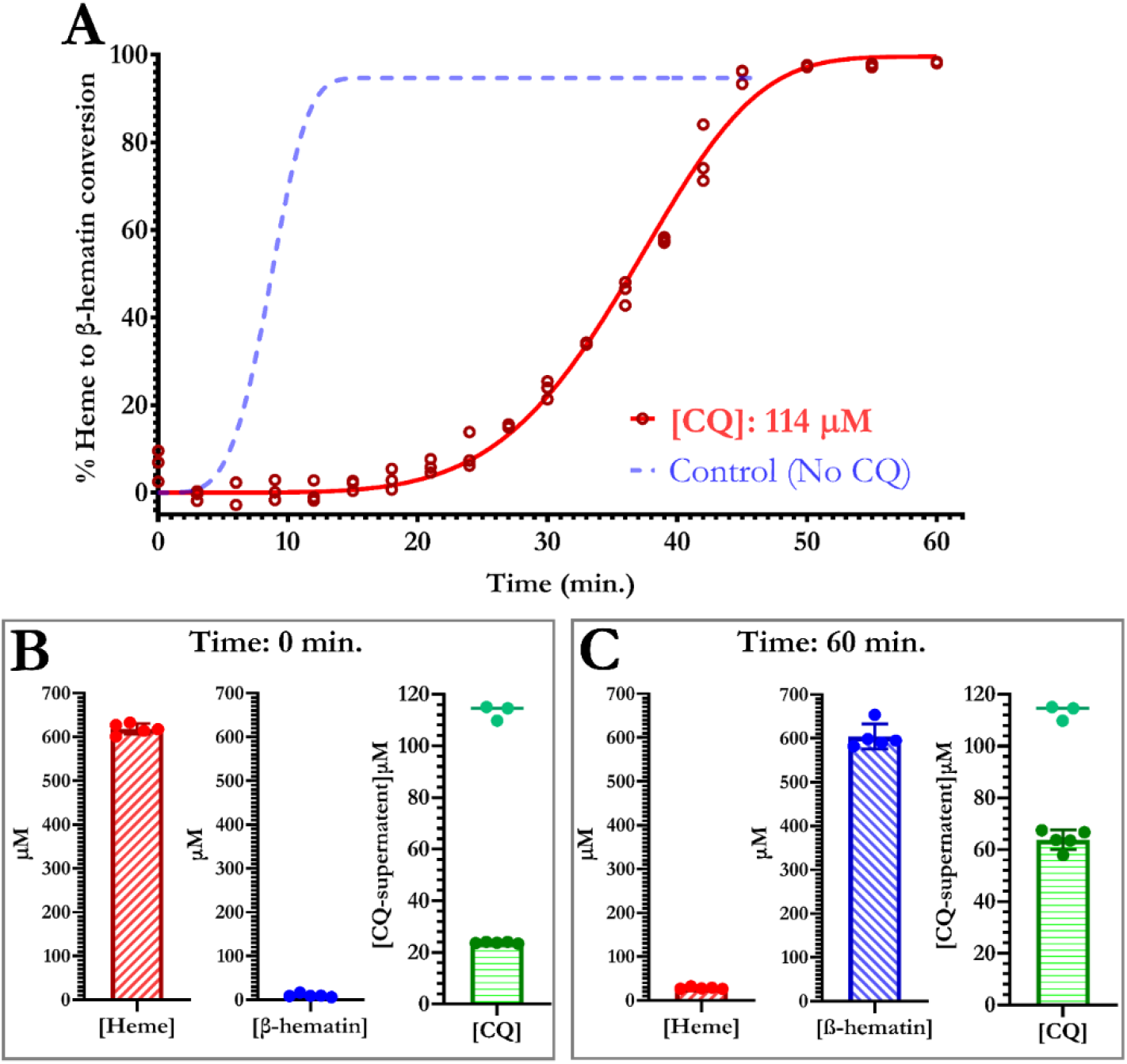
CQ pull-down assay: A, Time kinetics of heme to β-hematin transformation with a reaction consisting of 600 μM heme, 1.1 μM HDP, and 114 μM CQ (solid line). Each dot represents a replicate (N=3), and the solid line represents the best fit to the JMEK equation. The fitting statistics are given in Table S5. The dotted line (control) corresponds to the best fit to the JMEK equation of the data shown in Figure 2B, which represents the transformation series with 0 μM CQ. Inset B and C show the actual concentrations of heme substrate, β-hematin product, and CQ present in the supernatant at 0 and 60 min, respectively, from the onset of the reaction (A; solid red line; CQ: 114 μM). Free heme substrate is quantified via the pyridine hemochrome assay using a standardized curve (Fig. S3B). β-hematin product is quantified by dissolving the product in 0.1 N NaOH and measuring the solution’s absorbance at 385 nm (extinction coefficient of 58.4 mM^-1^ cm^-1^). Quantification of CQ is described in the materials and methods section, and the readings at the top (cyan) represent the quantity of CQ used in the transformation reaction (3A; 114 μM).

At the beginning of the transformation reaction (T=0), only 23.6± 0.26 μM out of 114 μM initially present CQ was available in the supernatant (Fig. 3B), indicating that around 87.4 μM of CQ had complexed with heme and precipitated. Interestingly, the pyridine hemochrome assay could accurately quantify the amount of untransformed heme present in the reaction mixture, even though it was in complex with CQ (Fig. 3B). This suggests that the acid-base interaction between heme and CQ in the heme-CQ complex can easily be disrupted by pyridine hemochromogen formation in the used assay. The transformation reaction reached completion in a characteristic sigmoid path (Fig. 3A), and the time required to complete the reaction was significantly larger in the reaction series with 114 μM CQ compared to the one without CQ (control).

At the end of the transformation reaction, only 27.37±2.18 μM of the initially present 617.8±1.11 μM heme substrate remained untransformed to β-hematin, resulting in 96% transformation (Fig. 3C). The β-hematin produced during the transformation reaction remained immune from getting decomposed by pyridine due to its crystalline nature (Fig. 3C). This β-hematin was dissolved into 0.1N NaOH, equivalent to the sampling volume (0.5ml) of time kinetics. Dissolved β-hematin is then quantified by measuring the solution’s absorbance at 385 nm (extinction coefficient of 58.4 mM^-1^.cm^-1^) and found to be 595.±25.54 μM (Fig. 3C). After the reaction, 61.78±1.11 μM of CQ was present in the supernatant, indicating that despite the completion of the transformation reaction, 52.2 μM of initially present 114 μM CQ was still complexed and present in the pellet. Since >95% of heme is transformed to β-hematin upon completion of the reaction, it is safe to assume that precipitated CQ is in complex with β-hematin as most of the heme substrate is consumed in the transformation reaction.

We repeated the experiment with 90 μM CQ to perturb the transformation of 600 μM heme into β-hematin, and the transformation followed the same sigmoid path to completion (Fig. S4). During this transformation reaction, the initially (T=0) precipitated 80 μM out of 90 CQ became free of complexation, raising the free CQ concentration in the supernatant from 10 μM to 46 μM as the transformation reaction reached completion. However, 50% of the initially available 90 μM CQ remained complexed with β-hematin upon completion of the reaction (Fig. S4).

### SEM and XRD

To visualize the influence of CQ on β-hematin crystal growth, we carried out heme to β-hematin transformation in reaction series with the same reaction constituents (600 μM heme, 0.098 μM HDP) but varying CQ concentrations ranging between 0 to 800 μM. In malaria parasites, the trophozoite stage, which is responsible for Hz production, lasts for around 12 hours [35]. Small quantities of HDP (0.098 μM) were used to ensure the time duration of 12 hours for transformation reaction completion in the reaction series without CQ (control). The range of CQ concentration for the experiment was selected by considering the property of quinolone bioaccumulation inside the *Plasmodium’s* digestive vacuole [30], where it could reach sub-millimolar concentrations. All reaction series, including one with an identical (600μM) and higher CQ concentration (800μM) for the used heme (600μM) substrate concentration, showed 98.4 % completion of heme to β-hematin transformation. Comparatively, the control reaction series without HDP showed no conversion of heme into β-hematin. The field emission scanning electron microscopy (FESEM) micrograph of β-hematin samples from the respective series showed a proportional increase in β-hematin-particle length (13.64 fold; 149.7±37 nm to 2042±764.9 nm) as well as width (9.68 fold; 51±8.9 nm to 494.1±144.1 nm) (Fig. 4A & S5). The dimensions of β-hematin crystal particles grown in the absence of CQ were comparable (149.7±37 nm; 51±8.9 nm) to native Hz crystals grown within the parasite (Fig. 4B), [36]. With increasing concentration of CQ in the reaction mixture, lamellar crystal growth was observed, suggesting unfinished growth (Fig. 4C). Similar growth patterns were earlier observed in native Hz crystals of *Plasmodium* gallinaceum [36] and validates that β-hematin crystal grows via classical layer mechanism [28].

**Figure 4:**
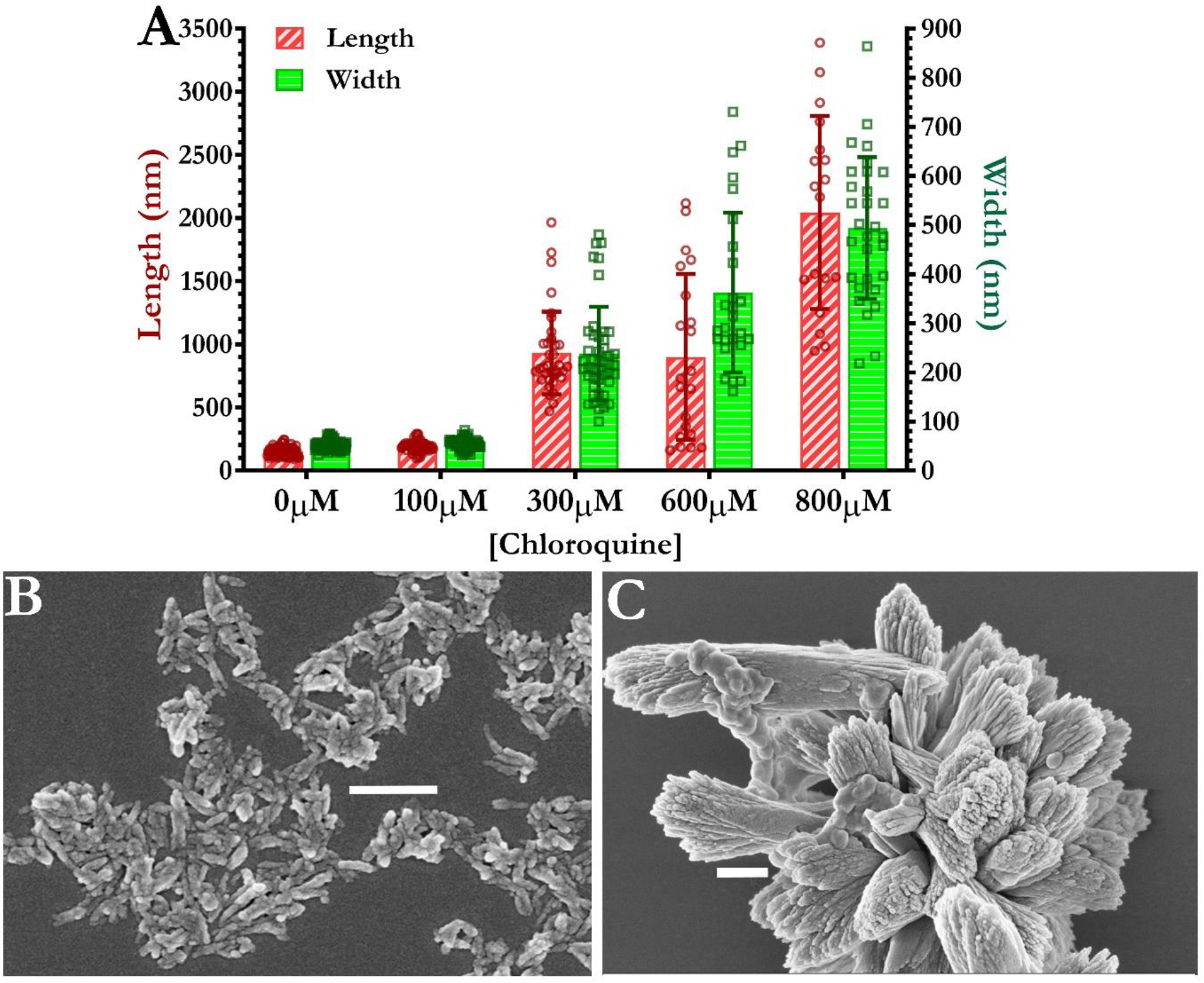
Influence of CQ on crystal growth: ***A***, Comparison of β-hematin crystal grain size (length on left Y-axis and width at right Y-axis; color-coded) grown at different concentrations of CQ. The error bar depicts standard error. Individual data points are overlaid with bar graphs. Electron micrograph of β-hematin crystals grown in the absence (***B***) and 800μM CQ presence (***C***). The electron micrograph is at different magnifications; therefore, the reference might be of varying size, but they correspond to a common length of 0.5 μm.

We performed XRD experiments to study the effect of CQ on the β-hematin crystallite size within the crystal particle and the overall structure. The samples of β-hematin used during this experiment were grown under the same experimental condition (Heme 600 μM and HDP 0.1 μM) except for the presence and absence of 800μM CQ during the transformation of heme into hemozoin transformation. The striking similarity between the XRD patterns (Fig. S6) and the unit cell parameters (Table S6) of both β-hematin samples shows that CQ, even at high concentrations, does not influence the β-hematin crystal structure. However, our XRD analysis shows that around three-fold increase in the crystallite size within the crystal particle (16.6±1 to 52.5±5 nm) in the sample of β-hematin grown in the presence of 800 μM CQ.

## DISCUSSION

The HDP-mediated rapid transformation of heme to β-hematin enables us to make several observations unique to this study. We selected the pH of 5.2 for our preliminary experiments since all earlier studies on HDP were performed on this pH [19][37][38][39][40][41]. However, most experiments were repeated at pH 4.8 to simulate better the condition within the parasite’s digestive vacuole, i.e., the Hz production site[42][43]. In addition, the selection of pH 4.8 was also influenced by the fact that pH of 4.8 has been widely used in earlier studies on the in vitro kinetics of heme to β-hematin [29][44][45][46][28]. The influence of CQ on heme to β-hematin transformation was studied by inhibition and time-kinetics experiments. The time-kinetic data were analyzed via the KJMA equation [31]-[32], commonly used to study isothermal phase transformation reactions. This analysis always yielded sigmoid curves, characteristic of phase transformation during crystallization (Fig. 1A, 2 & S2) [47]. Three stages characterize this sigmoid curve: nucleation, crystal growth, and stationary (Fig. 5A). The nucleation stage marks the beginning of the time-kinetic reaction, where the best-fit line behaves asymptotically [47]-[48]. Crystal nucleation events predominate this stage, leading to a minuscule transformed phase. The crystal-growth stage follows the nucleation stage, where crystal nuclei grow in size at a constant velocity, leading to an almost linear appearance of the best-fit line. Finally, due to the impingement of crystals on each other or the limitation of the substrate, the best-fit line again becomes asymptotic, marking the saturation stage.

**Figure 5.**
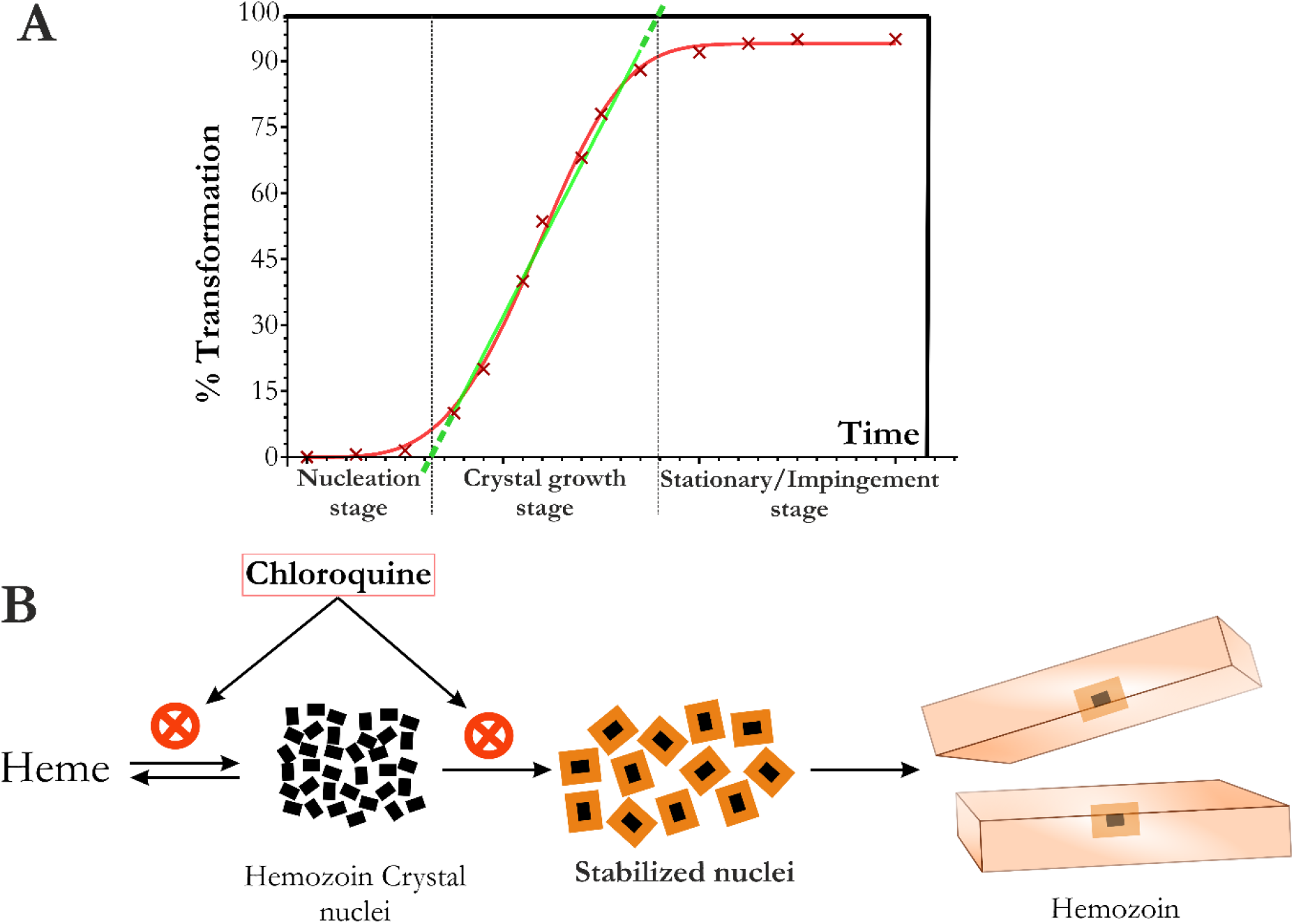
Phase transformation during heme to Hz conversion. ***A***, The sigmoid curve of phase transformation kinetics is commonly interpreted into three stages, nucleation, crystal growth, and saturation stages. The X-axis intercept of the linear crystal growth phase (green line) is a good approximation for both the nucleation and saturation stages. ***B***, The schematic of a model displaying CQ’s mode of action.

Antimalarial derived from quinine, including CQ, induce toxicity in *Plasmodium* by interfering with Hz production [23][24][25][26][27][28][29][30]. Earlier studies suggest that quinoline-derived antimalarials form non-crystallizable complexes with free heme [49][50]. It is further believed that this CQ-heme complex is also cytotoxic and responsible for the malaria parasite’s demise [51]. As a result of this complex formation, heme availability for Hz production decreases, thereby inhibiting the transformation process [23][24][25][26]. Therefore it is expected that in the presence of CQ, the percent transformation of heme into β-hematin must be smaller than the control reaction (without CQ). However, our time-kinetics experiments show that the presence of CQ in the reaction mixture does not affect the final amount of phase transformed (Fig. 2B, 3 & S2B). Our CQ pull-down assay shows that acid-base interaction in the heme-CQ complex can easily be perturbed to make heme available for the β-hematin formation (Fig. 3). Therefore, the complexation between heme and CQ does not compromise the ability of heme to transform into β-hematin. Further, against the general convention, our studies show that CQ does not stop but slows down the transformation of heme into β-hematin. This observation is true even when CQ is in molar excess for heme (Fig. 4). Therefore, the CQ IC50 values during heme to β-hematin transformation are not fixed but variable depending upon both heme concentration and observation time (Fig. 1 & S1).

Our CQ pull-down assay shows that, similar to heme, β-hematin can also bind to CQ (Fig. 3B). However, the XRD data clearly shows that the crystal structure of the β-hematin sample grown in the presence and absence of CQ are similar. Therefore, CQ precipitated out from the solution via β-hematin is likely to adhere on the surface of the β-hematin crystals and does not perturb the internal crystalline arrangement. This observation agrees with previously reported studies has suggested that the quinolone family of drugs, primarily CQ binds to the Hz crystal surface and inhibits further substrate addition resulting in the poisoning of Hz crystal growth [27][28][29][30]. Therefore it is expected that in the presence of CQ, the growth of β-hematin crystal should either stop or decreases significantly. In contrast, our SEM and XRD data clearly show that the size of overall crystal particles and crystallite, respectively, increases in the presence of CQ. In addition, arguably, in many of the earlier studies where CQ is shown to inhibit Hz crystal growth [30], the size of Hz crystals is arguably comparable to wild-type native crystals [36].

Overall, our data clearly shows that the CQ does form complex with free-heme (and β-hematin) but without compromising its capacity to transform into β-hematin. However, CQ does increase the time duration to complete heme to β-hematin transformation but in a way that results in larger crystals particle and crystallite size. The nucleation event of the β-hematin crystals is probably the aspect of heme to β-hematin transformation left to be influenced by CQ and explains our observations. To explore this possibility, we performed a time-kinetics experiment in reaction series where CQ in a fixed concentration was introduced into the reaction mixture but at different time points during the time-kinetics experiments (Fig. 2C & S2C). Consistent with other time-kinetics experiments involving CQ, we observed that CQ increases the time required to complete the transformation reaction and simultaneously increases the nucleation stage duration. This influence of CQ on the transformation reaction is most evident when CQ is introduced at the very beginning of the transformation reaction, i.e., when the nucleation events predominantly populate the reaction. Therefore, as we increase the duration of the time-point to introduce CQ into the reaction mixture, this influence of CQ on the transformation reaction gradually decreases and eventually becomes comparable to the control reaction, suggesting that CQ interferes with the nucleation events of β-hematin formation. The CQ-induced smaller population of formed nuclei is expected to yield fewer but bigger β-hematin crystals due to non-limiting substrate (heme) availability, explaining our SEM and XRD data. In addition, if the CQ complexation-induced decreased free-heme concentration is vital to inhibit Hz formation, first and foremost, the nucleation events are likely to be inhibited, as nucleation is most sensitive to the active concentration of substrate. Similarly, if the coating of CQ on the Hz growth surfaces is required to inhibit crystal growth, smaller Hz crystal nuclei with a much smaller surface area should be more sensitive to this inhibitory phenomenon [52].

The freshly formed crystal nuclei are generally considered unstable [53]-[54] and in equilibrium with a supersaturated environment. Therefore, these nuclei evolve and dissolve until they grow to a critical size [53]-[54] (stabilized nuclei) to sustain the crystal growth (Fig. 5B). We believe that CQ either interferes with the formation of β-hematin nuclei or it may prevent the nuclei from reaching a critical size to form stabilized nuclei. However, once the crystal nuclei achieve the critical mass, they grow in size at a constant rate irrespective of the presence of CQ (Fig. 5B), explaining the completion of the transformation reaction even in the presence of CQ. We also believe that an antimalarial drug-like, like CQ, does not need to entirely arrest the Hz production process to induce toxicity in *Plasmodium*. Instead, the heme to Hz transformation can be slowed down to buy time for the accumulated heme to induce cytotoxicity in the parasite.

Our findings indicate that in the presence of CQ, the Hz crystal within Plasmodium parasites can grow to a size that may damage cellular compartments. Previous studies have shown that the phagocytosis of crystalline material by phagocytes can cause lysosomal damage [55][56][57][58], leading to the leakage of proteases into the cytosol, which can digest vital proteins and organelles like mitochondria. Such digestion can cause mitochondrial outer membrane permeabilization-dependent cell death [59]. The extent of lysosomal damage can determine whether cells undergo necrosis or activate a caspase activation cascade [59][60]. As the Plasmodium digestive vacuole is comparable to lysosomes in other eukaryotic cells, we suggest that large Hz crystals in the presence of CQ may compromise the integrity of the digestive vacuole of Plasmodium. This can cause the contents of the digestive vacuole to leak into the cytosol, leading to crystal-mediated toxicity. Although supported by previous studies in phagocytes, this hypothesis requires experimental validation before it can be considered valid.

This study considers the HDP as one of the proposed native mediators for heme to Hz transformation, as reported in previous literature [19][21][40][37][39][41][38][61]. However, it is possible that other mediators such as free lipids or lipids in the plasma membrane could actually be facilitateing this transformation within the parasite’s digestive vacuole [62]. Nonetheless, in our in-vitro settings, HDP was found to be the most efficient mediator of heme to β-hematin transformation [19], which allowed us to observe results that other studies may have missed. Since our study primarily focused on the effect of chloroquine on hemozoin formation, and it is well-established that chloroquine inhibits hemozoin production by interacting with either free heme [23][24][25][26][49][50][51] or hemozoin [27][28][29][30], rather than the catalyst involved in the transformation reaction, the majority of the findings presented in this study would hold true regardless of the mediator used. As with any catalyst, the HDP only affects the reaction’s cross-section without altering the nature of the reactants.

Although this study attempted to study the heme to β-hematin transformation under conditions close to the native state, it is still a limited representation of the in-vivo process inside the parasite. Nevertheless, the unique findings in this study should not be disregarded merely because of its reductionist nature. The insights obtained from a simple in-vitro system can, to some degree, translate to the processes occurring within the cell. Unfortunately, it is impossible to extract crystal nuclei from their surroundings to purify them since they are metastable entities that are too tiny to be seen using any known techniques. As a result, studying the impact of chloroquine on Hz nucleation in vivo may be unfeasible, and at best, similar to this study, it can only be studied in an in-vitro system.

## Conclusion

In this study, we studied HDP-mediated phase transformation of heme into β-hematin under CQ’s influence. This study suggests that CQ neither decreases the active concentration of free heme via complex formation nor interferes with the growth of Hz crystals to exhibit its antimalarial properties. In addition, Hz production is not stopped but slowed down by CQ via perturbing formation and stability of Hz crystal nuclei for the tested range of CQ concentrations. Therefore, we propose that CQ induces antimalarial properties by poisoning the nucleation event or nuclei stability of Hz crystal.

## EXPERIMENTAL PROCEDURES

### Material

Dimethyl sulfoxide (DMSO), hemin (Sigma-Aldrich, ≥97%, 51280-1G), sodium acetate, N-lauroyl Sarcosine (8147150100) and Tris base were obtained from Sigma-Aldrich. Pyridine, acetone and glacial acetic acid were procured from EMPURA, HEPES was procured from Himedia (MB016). Chloroquine diphosphate salt was procured from MP biochemicals (CAT No.: 193919)

### Methods

#### 1. Cloning, expression and purification

Cloning, expression and purification of HDP protein was performed as earlier explained [19][63].

#### 2. Enzymatic Assay

##### 2.1. Substrate preparation

The hemin solution was freshly prepared in 100% (v/v) DMSO (in dark) by vigorous shaking for 10 min followed by centrifugation at 17000g RCF for 10 min. The supernatant was filtered through a glass microfiber filter (Whatman^®^ Cat no 1822 025) syringe filter to remove undissolved hemin crystals. The concentration of hemin stock was determined via UV-vis absorption spectroscopy via diluting the stock in 0.1 N NaOH and measuring absorbance at 385 nm (extinction coefficient, 58.4 mM^-1^ cm^-1^). The solution was used within 1 hour. Similarly the chloroquine stock was prepared in 100mM sodium acetate buffer pH 4.8. The concentration of chloroquine was also measured UV-vis absorption spectroscopy by measuring the absorbance at 329 nm and using an extinction coefficient of 16.6 mM^-1^ cm^-1^ (British Pharmacopoeia, HMSO, London, GB: 1980, page. 103)

##### 2.2. Assay

The pyridine hemochrome assay was performed as previously described (37). All assay was performed at pH 5.2 and 4.8 at 37 °C in an assay buffer containing 0.25 M sodium acetate and 5% (v/v) DMSO. HDP was used in the concentration range of 0.01 to 0.5 μM. After incubation, the reaction was stopped by changing the pH of the reaction buffer from 5.2 to 7.5 via the addition of a stock solution of 3M Tris-Cl pH 7.5. The unconsumed heme substrate left in the reaction mixture was detected by adding pyridine reagent (40% v/v pyridine, 20% v/v acetone, 200 mM HEPES pH 7.5) to the reaction mixture, to achieve the final pyridine concentration of exactly 5% (v/v). The β-hematin product remain insoluble and was removed out of reaction mixture via centrifugation. The quantification of free heme was done by measuring the absorbance at 405 nm via UV-vis absorption spectroscopy, within 10-30 minutes of adding pyridine reagent. For concentrated solution, dilution with a base solution containing 4% v/v pyridine, 2% v/v acetone and 20 mM HEPES pH 7.5. The amount of heme to β-hematin transformation in a reaction was assessed by measuring the amount of heme substrate consumed during the reaction in respect to control reaction (reaction without HDP). That is, in a reaction the amount of heme substrate consumed is equal to amount of β-hematin produced. All measurements were made in cuvette of 2 mm path length via a double-beam spectrophotometer (Jasco-V630).

###### Note

For 12 hours at 37°C, without HDP, a negligible amount of heme substrate transform into β-hematin at pH 4.8 and 5.2 (data not shown).

##### 2.3. Inhibition studies

The common assay procedure used in the inhibition studies is described in section 2.2 of material and methods. The inhibition studies were performed at pH of 5.2 and 4.8 in series with different concentrations of heme substrate among series. Each series constituted of reaction of equal volume and heme substrate concentration but different CQ concentrations. The reactions in each series were stopped at a specified time point. The production of β-hematin was studied as a function of increasing CQ concentration. The reaction in each series was stopped at specified time point. The amount of β-hematin produced in reaction was measured by assaying free heme only. Normalization of data was done for each series separately, with a 100% value corresponding to the amount of β-hematin produced without inhibitor CQ and nil as 0%. Half the maximal inhibitory concentration (IC50) value was deduced by fitting the data to the following equation (Equation-1) using least-squares fitting methods with GraphPad Prism v5.0 (USA).

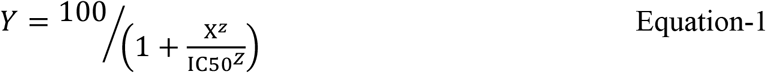

Where “Y” is normalized (%) β-hematin production, “X” chloroquine concentration in μM and “Z” is Hill slope.

##### 2.4. Time-Kinetics

The common assay procedure used in the inhibition studies is described in section 2.2 of material and methods. Time-kinetics experiments were performed at 37 °C in a reaction volume of 15 to 50 ml at pH 5.2 and 4.8 with constant constituents (substrate, HDP and/or chloroquine). Aliquots of 0.3 to 0.5 ml were removed from the reaction suspension at different time intervals in a 1.5 ml micro centrifuge tube prefilled with 3M Tris-Cl pH 8.5 for changing the pH of the reaction mixture to 7.5 and instantaneously stopping the reaction. Two aliquots were removed simultaneously from the same reaction suspension (N=1) or two independent reactions (N=2) for each time interval. Normalization of data was done with respect to 100% value corresponding to β-hematin concentration equivalent to amount of un-transformed heme available at the beginning of the reaction (T=0 min.) and nil as 0 %. Time-kinetics of β-hematin formation was generally analyzed by fitting the data into Kolmogorov-Johnson-Mehl-Avrami (KJMA) equation (Equation-2) using least-squares fitting with GraphPad Prism v5.0 (USA).

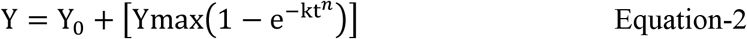

Where “Y” is the fraction of substrate converted to β-hematin in %, “Y_0_” is the fraction of substrate converted at zero minutes, “Ymax” is the maximum percentage of β-hematin formed at the end of the reaction, “k” is an overall kinetic rate constant, “t” is time in min., and “*n*” is the Avrami constant.

In some case the data was analyzed by fitting the data into biphasic sigmoid growth curve equation (Equation-3) using least-squares fitting with Origin (USA)

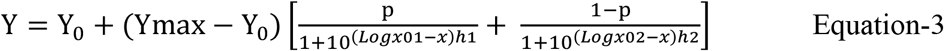

Where “Y” is the fraction of substrate converted to β-hematin in %, “Y_0_” is the fraction of substrate converted at zero minutes, “Ymax” is the maximum percentage of β-hematin formed at the end of the reaction, “h” (1 & 2) is slope and “p” is proportion.

#### 3. Field Emission Scanning Electron Microscopy and X-ray diffraction powder diffraction

##### 3.1. Sample preparation

The β-hematin sample of Field Emission Scanning Electron Microscopy (FESEM) and powder X-ray powder diffraction (XRD) measurements were prepared in six reaction series, each with a 30 ml reaction volume. All series have the same common reaction condition of 500mM NaAc pH 5.2, 600 μM Heme, and 0.1 μM HDP, except varying CQ concentrations (0, 100, 300, 600, and 800 μM). One of the six reaction series was a negative-control reaction series without CQ or HDP. In all reaction series, chloroquine was introduced at the beginning of the reaction mixture preparation, with HDP being added last. The reaction was carried out in non-shaking conditions at 37°C for 7 to 10 days. The reaction volume of 4ml was removed from each reaction series was centrifuged to get the β-hematin pellet. The pellet from each reaction series was washed repeatedly, with each washing step involving harvesting the pellet by centrifugation. The washing steps sequentially involve washing twice with MQ followed by washing with 5% pyridine twice, once washing with 100 % (v/v) DMSO, and finally twice washing with MQ. For XRD measurements, the washed pellet from the respective reaction series was again washed with 100 % (v/v) ethanol, followed by drying the centrifuged pellet on RT. For FESEM studies, the washed pellet is re-suspended in 400 μl of MQ and sonicated for half an hour. Volume 4 μl of the sonicated sample was then placed on a glass slide and dried under a mild vacuum. A conductive chromium coating via sputtering was placed over a glass slide with deposited samples.

##### 3.2. Electron microscopy

The effect of CQ on the surface morphology of β-hematin was studied by FESEM (Sigma02, Carl Zeiss). ImageJ software performed the particle size analysis on the collected electron micrograph images.

##### 3.2. XRD

The XRD data were collected at 25°C on Extreme Conditions Angle Dispersive/Energy dispersive X-ray diffraction beamline (BL-11) at the Indus-2 synchrotron source. All measurements were carried out at 12 keV (1.0232 Å) with the sample mounted between the scotch tape. The collected data was refined for background, lattice parameters, and crystallite size via Rietveld Refinement using GSAS II Software. The structure coordinates of earlier submitted β-hematin (CCDC deposition no. 841403) were used as a reference during the refinement. The XRD data of NIST RSM 674b (CeO2) was used during refinement to account for peak broadening due to instrumentation. The refined unit cell parameters and crystallite size is described in **table 1**.

#### 4. CQ pull-down assay

CQ and heme in defined concentrations are agitated extensively (vigorous mixing, on a vortex machine) to make them interact in an acidic pH 4.8 (Sodium acetate 0.5M). Heme is insoluble in reaction buffer at acidic pH, whereas chloroquine is reasonably soluble. The agitated suspension is centrifuged at 16000g (RCF) for 30 min to pellet down the insoluble heme. During this centrifugation, the CQ-Heme complex also gets precipitated, leaving uninteresting CQ in the supernatant. This uninteresting CQ is then quantified by measuring absorbance at 329 nm using an extinction coefficient of 16.6 mM^-1^ cm^-1^ (British Pharmacopoeia, HMSO, London, GB: 1980, page. 103). The CQ in the CQ-heme complex is measured by subtracting the amount of uninteresting CQ from the total amount of CQ initially added to the reaction.

##### Note

The amount of CQ predicated with heme extensively depends on and is proportional to the agitation. Aliquots out of the same sample with different agitation times tend to have a different amount of CQ precipitation. Therefore this method is not ideal to estimate the stoichiometry of heme-CQ complex.

## Supporting information

This article contains supporting information. The supporting information consisted of four figures (Fig. S1 to S6) and six tables (Table S1 to S6)

## DATA AVAILABILITY

All data are contained within the manuscript.

## ACKNOWLEDGMENTS

We acknowledge the use of the biochemical user-lab facility of Px-Bl21 and Extreme Conditions Angle Dispersive/Energy dispersive X-ray diffraction beamline (BL-11) of Indus2 synchrotron source. Special thanks to Dr. Pragya Tiwari for our technical discussions and SEM sample preparation. We thank Shri Rakesh Kaul for providing the FESEM facility.

## AUTHOR CONTRIBUTIONS

Rahul Singh: Conceptualization, Methodology, Experimentation, Analysis, Writing-Original draft preparation; Rashmi Singh: FESEM images acquisition; Velaga Srihari: XRD data acquisition and refinement; Ravindra D. Makde.: Supervision, Analysis, Writing-Reviewing.

## COMPETING INTEREST STATEMENT

All authors declare no conflicts of interest with the contents and findings of this article.

## ABBREVIATIONS AND NOMENCLATURE

HDP: heme detoxification protein
Hz: hemazoin
CQ: chloroquine
IC50: Half-maximal inhibitory concentration
KJMA: Kolmogorov-Johnson-Mehl-Avrami
n: Avrami constant

## Notes

### Competing Interest Statement

The authors have declared no competing interest.

