## Supplementary material for "Revealing Chloroquine’s Antimalarial Mechanism: The Suppression of Nucleation Events during Heme to Hemozoin Transformation": This article contains supporting information. The supporting information consisted of four figures (Fig. S1 to S6) and six tables (Table S1 to S6)

RESULTS: Supplementary figures and tables:

**Table S1:** CQ mediated Inhibition (Figure 1).

| **Parameters** | **5 Min.** | | | | **7 Min.** | | | |
|---|---|---|---|---|---|---|---|---|
| | **200 µM** | **330 µM** | **460 µM** | **600 µM** | **200 µM** | **330 µM** | **460 µM** | **600 µM** |
| [HDP]µM | **1.5** µM (constant) | | | | | | | |
| pH | **4.8** | | | | | | | |
| IC50 (µM) | 10.96±1.06 | 18.6±1.11 | 35.86±1.03 | 41.68±1.05 | 18.66±1.02 | 24.23±1.072 | 42.2±1.04 | 45.58±1.04 |
| $R^2$ | 0.9502 | 0.9031 | 0.9893 | 0.9702 | 0.9966 | 0.9529 | 0.9763 | 0.9724 |
| Sy.x (a) | 7.605 (24) | 12.68 (24) | 3.839 (24) | 6.111 (24) | 2.149 (24) | 8.985 (24) | 5.858 (24) | 6.043 (24) |

The standard deviation of the residuals (Sy.x) = $\sqrt{\frac{\Sigma(residuals^2)}{a-K}}$ where "*a*" is the number of data points, "*K*" number of parameters fit by regression and residual is the vertical distance (in Y units) of the point from the fit line.

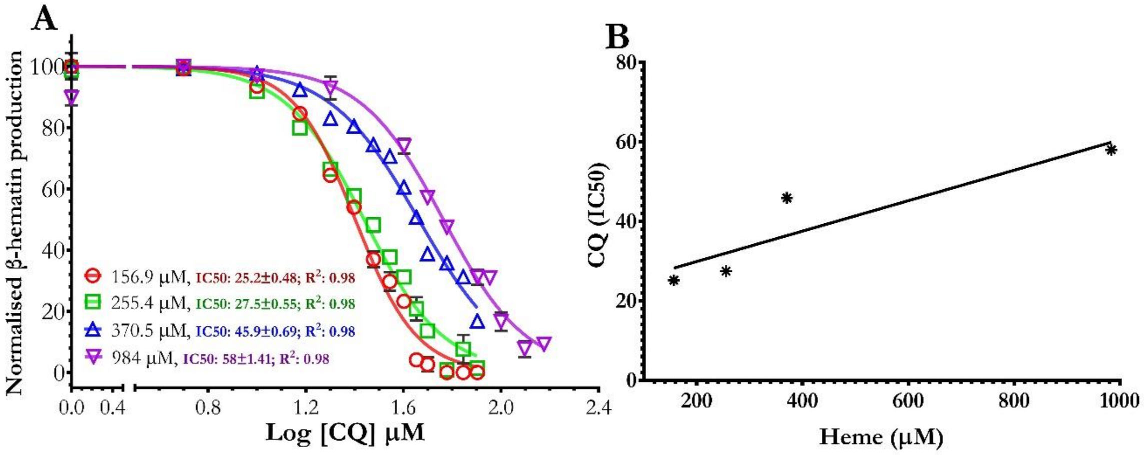


Figure S1: *Chloroquine mediated inhibition of β-hematin production (pH5.2). A,* The graph represents a change in normalized β-hematin production as a function of chloroquine concentration at the constant HDP and color-coded heme concentrations. Individual points (square, circle, or triangle) represent the mean value calculated by measuring the change in free-heme concentration present in assay supernatant. The bar represents the error between replicate reactions (N=3). The fitting statistics is given in the Table S2. *B,* The graph shows a positive trend between the reaction's CQ IC50 value and heme concentration.

**Table S2:** Chloroquine Inhibition Assay (Figure S1).

| Parameters | Heme concentration | | | |
|---|---|---|---|---|
| | **156.9 μM** | **255.4 μM** | **370.5 μM** | **984 μM** |
| **pH** | 5.2 | | | |
| *Z* (Hill Slope) | 3.43±0.2 | 2.69±0.14 | 2.38±0.10 | 2.55 ± 0.16 |
| IC50 (μM) | 25.25±0.48 | 27.46±0.55 | 45.93±0.69 | 58.00 ± 1.41 |
| $R^2$ | 0.987 | 0.986 | 0.986 | 0.984 |
| Sy.x (a) | 4.510 (28) | 4.300 (28) | 3.299 (28) | 4.497 (28) |

The standard deviation of the residuals (Sy.x) = $\sqrt{\frac{\sum(\boldsymbol{residuals^2})}{\boldsymbol{a-K}}}$ where "*a*" is the number of data points, "*K*" number of parameters fit by regression and residual is the vertical distance (in Y units) of the point from the fit line.

**Table S3:** Curve fitting statistics of Figure 2; time-kinetics at pH 4.8.

| **Figure 2A; time-kinetics at different HDP concentrations** | | | |
|---|---|---|---|
| Parameters | HDP Concentration | | |
| | **0.75** µM | **1** µM | **1.5** µM |
| [Heme] µM | 600 | | |
| pH | 4.8 | | |
| Fitting Kolmogorov-Johnson-Mehl-Avrami equation | | | |
| n | 5.88±0.22 | 4.246±0.02 | 3.938±0.22 |
| Ymax | 94.68±0.787 | 94.5±0.61 | 87.80±0.6209 |
| $R^2$ | 0.997 | 0.994 | 0.996 |
| Sy.x (a) | 2.179 (24) | 2.969 (34) | 2.171(24) |

| **Figure 2 *B*; time-kinetics at different CQ concentrations.** | | | | |
|---|---|---|---|---|
| Parameters | [CQ] (µM) | | | |
| | 0 µM | 50 µM | 75 µM | 100 µM |
| [HDP] µM | 1 | | | |
| [Heme] µM | 600 | | | |
| pH | 4.8 | | | |
| Fitting Kolmogorov-Johnson-Mehl-Avrami equation | | | | |
| Ymax | 94.7±0.51 | 94.72± 0.71 | 95.27± 1.07 | 97.02± 2.4 |
| n | 4.22±0.02 | 3.63±0.01 | 5.28±0.29 | 3.65±0.18 |
| $R^2$ | 0.9947 | 0.9930 | 0.9926 | 0.9910 |
| Sy.x (*a*) | 2.76 (40) | 3.45 (40) | 3.65 (40) | 3.42 (40) |

| **Figure 2C; Influence of CQ introduction at different time points on the kinetics of heme to β-hematin conversion.** | | | | | |
|---|---|---|---|---|---|
| **Parameters** | **Control** | **A** | **B** | **C** | **D** |
| [HDP] µM | 0.8 Constant) | | | | |
| [Heme] µM | 600 (Constant) | | | | |
| pH | 4.8 | | | | |
| [CQ] µM | Not applicable | 100 (Constant) | | | |
| CQ introduction time (min.) | | 0 | 6 | 11.5 | 17.5 |
| **Fitting** | KJMA (Eq.2) | | Biphasic sigmoid (Eq.3) | | KJMA (Eq.2) |
| Ymax | 99.01 | 99.13 | 100.4 | 100.2 | 98.45 |
| n | 3.385 | 5.349 | Not applicable | | 1.984 |
| $R^2$ | 0.9944 | 0.9936 | 0.9987 | 0.99 | 0.9736 |
| Sy.x (*a*) | 2.431 (58) | 2.981 (58) | *4.177* (58) | *4.526* (58) | 5.213 (58) |

The standard deviation of the residuals (Sy.x) = $\sqrt{\frac{\Sigma(residuals^2)}{a-K}}$ where "a" is the number of data points, "K" number of parameters fit by regression and residual is the vertical distance (in Y units) of the point from the fit line.

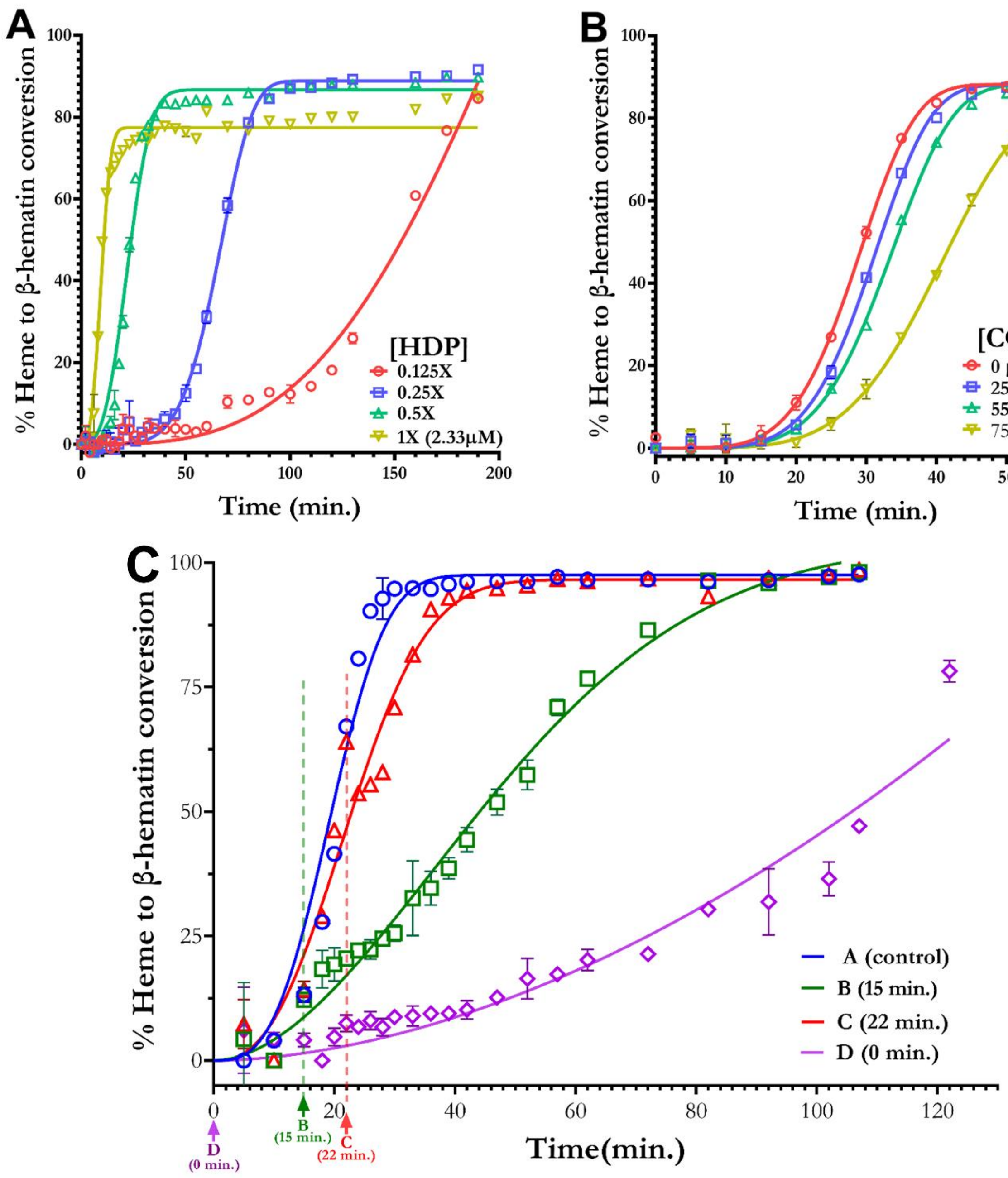


Figure S2. *Time-kinetics under the influence of CQ*. *A*, The graph shows the change in a percentage of heme transformed to β-hematin as a function of time at constant substrate (880 µM) concentration and color-coded series with different HDP concentration. *B*, Graph shows % heme to β-hematin transformation as a function of time at a constant concentration of HDP (0.32 µM) and heme substrate (984 µM), but different chloroquine concentrations (color-coded) in each reaction series. The bar represents the error between duplicates reactions (N=1) *C*, The graphs represent a change in normalized β-hematin production as a function of time at the constant HDP (0.75 µM) and heme (600 µM) concentrations; each reaction series except control series (seriesA) was exposed to 100 µM chloroquine at a different time point after the onset of reaction, i.e., 0 min. (series D), 15 min. (series B), and 22 min. (series C) represented by color-coded dotted lines and arrows. The line represents the best fit to the KJMA equation (color-coded), the individual points (square, circle, or triangle) represent the mean value calculated by measuring the change in free-heme concentration present in assay supernatant. The bar represents the error between replicate reactions (N=2). The fitting statistics is given in the Table S4.

**Table S4:** Curve fitting statistics of Figure S2

| **Figure S2A; time-kinetics at different HDP concentrations** | | | | |
|---|---|---|---|---|
| Parameters | HDP (1X=2.33 µM) | | | |
| | **0.125X** | **0.25X** | **0.5X** | **1X** |
| [Heme] µM | 880 | | | |
| pH | 5.2 | | | |
| Fitting Kolmogorov-Johnson-Mehl-Avrami equation | | | | |
| n | 3.23±0.33 | 5.47±0.20 | 2.91±0.01 | 2.93±0.22 |
| Ymax | 228.5±164 | 88.8±0.60 | 86.62±0.84 | 77.38±0.58 |
| $R^2$ | 0.976 | 0.996 | 0.987 | 0.979 |
| Sy.x (a) | 3.38 (62) | 2.355 (62) | 4.391(62) | 3.845 (62) |
| Figure S2 *B*; time-kinetics at different CQ concentrations. | | | | |
| Parameters | [Chloroquine] (µM) | | | |
| | **0 µM** | **25 µM** | **55 µM** | **75 µM** |
| [HDP] µM | 0.36 | | | |
| [Heme] µM | 984 | | | |
| pH | 5.2 | | | |
| Fitting Kolmogorov-Johnson-Mehl-Avrami equation | | | | |
| Ymax | 88.12±0.47 | 88.03± 0.53 | 88.15± 0.14 | 84.19± 1.23 |
| n | 4.73±0.13 | 5.07±0.15 | 4.97±0.14 | 4.686±0.17 |
| $R^2$ | 0.999 | 0.999 | 0.999 | 0.997 |
| Sy.x (*a*) | 1.36 (26) | 1.47 (26) | 1.41 (26) | 1.697 (26) |
| **Figure S2C; Influence of CQ introduction at different time points on the kinetics of heme to β-hematin conversion.** | | | | |
| **Parameters** | **A Control** | **B** | **C** | **D** |
| [HDP] µM | 0.75 (Constant) | | | |
| [Heme] µM | 600 (Constant) | | | |
| pH | 4.8 | | | |
| [CQ] µM | Not applicable | 100 (Constant) | | |
| CQ introduction time (min.) | | 15 | 22 | 0 |
| Fitting Kolmogorov-Johnson-Mehl-Avrami equation | | | | |
| Ymax | 97.57±1.476 | 104.1±3.18 | 96.61±1.76 | Ambiguous |
| n | 3.04±0.024 | 1.86±0.09 | 2.53±0.182 | |
| $R^2$ | 0.9671 | 0.9839 | 0.9688 | |
| Sy.x (*a*) | 6.149 (46) | 4.184 (46) | 5.747 (46) | |

The standard deviation of the residuals (Sy.x) = $\sqrt{\frac{\sum(residuals^2)}{a-K}}$ where "a" is the number of data points, "K" number of parameters fit by regression and residual is the vertical distance (in Y units) of the point from the fit line.

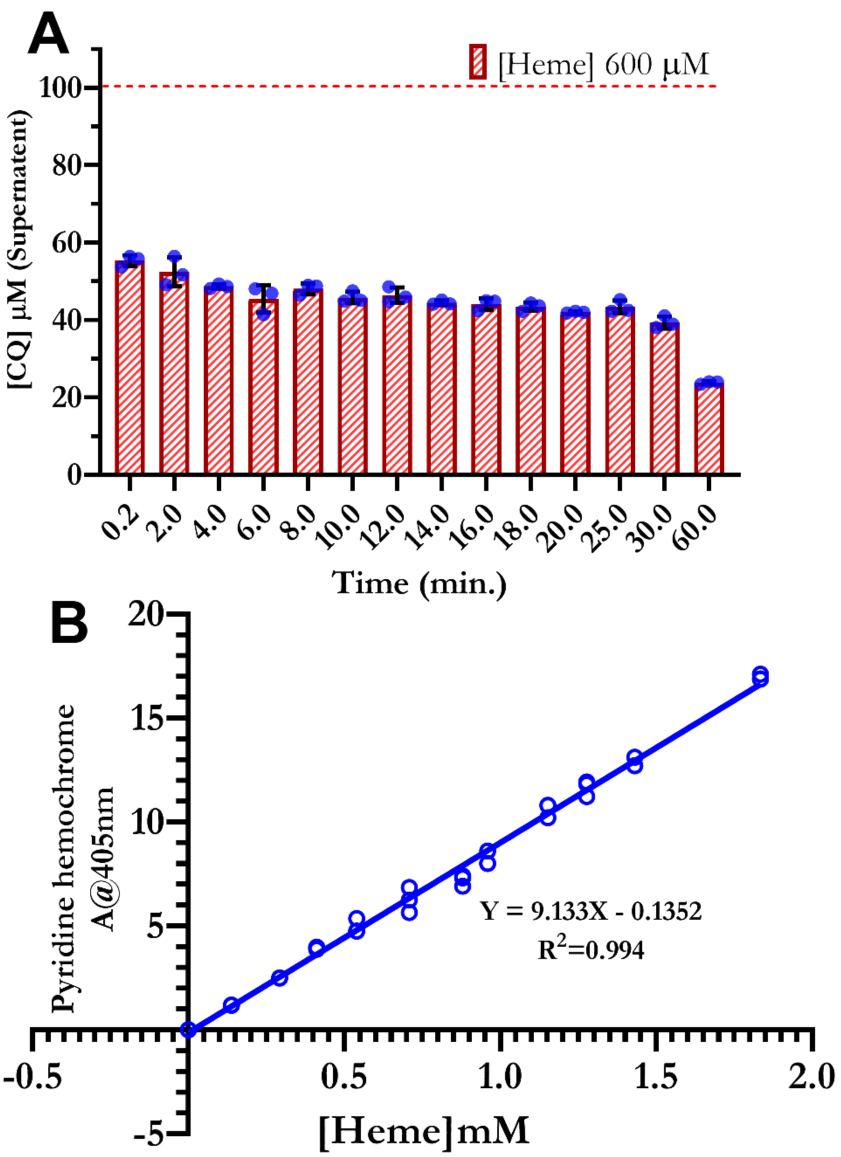


**Figure S3:** *Chloroquine precipitation with mixing time*: **A,** Reaction mixture (without HDP) with 600 µM heme and 100 µM CQ (dotted line) was vigorously mixed for a variable time. The bar represents the concentration of CQ left in the supernatant for the corresponding mixing time. Each dot represents a replicate (N=3) and the error bar represents the error between replicates. **B**, Beer's law plot at 405 nm with 5% (v/v) aqueous pyridine.

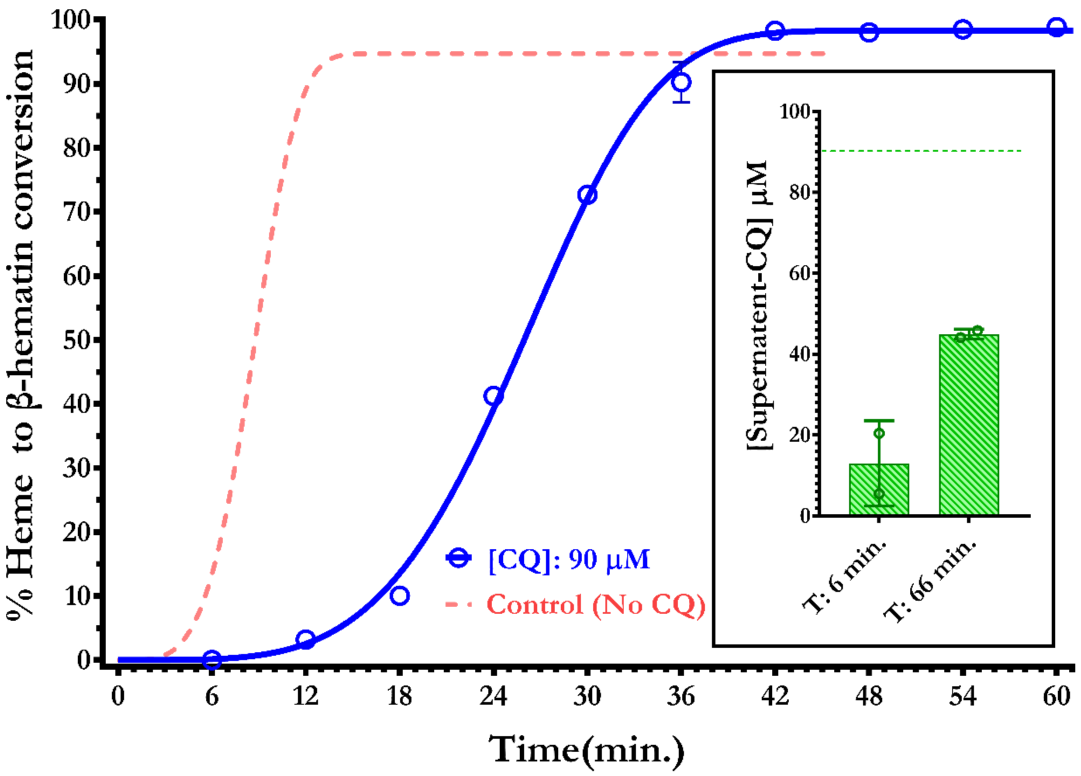


Figure S4: *CQ pull-down assay:* Time kinetics of heme to β-hematin transformation with reaction consisting of 600 μM, 1.1 μM HDP, and 90 μM CQ (solid line). The dotted line (control) corresponds to the best fit to the JMEK equation of the data shown in Figure 2B, representing the transformation series with 0 μM CQ. The error bar represents the error between replicates (N=2). The fitting statistics are given in Table S5. The inset shows the actual concentrations of CQ present in the supernatant at the beginning (T=6 min.) and completion (T=60 min.), of the transformation reaction with respect to the 90 μM CQ (dotted line) added to the reaction.

**Table S5.** Curve fitting statistics of Figure S3B

| **Figure S3B; time-kinetics and CQ precipitation** | | | |
|---|---|---|---|
| [HDP] μM | 1.1 | | |
| [Heme] μM | 600 | | |
| pH | 4.8 | | |
| Fitting | Fitting Kolmogorov-Johnson-Mehl-Avrami equation | | |
| Figure | Figure S4 | Figure 3A | Figure 2 B; CQ 0μM |
| [CQ] μM | 90 | 114 | NO CQ (Control) |
| n | 4.28 | 5.282 | 4.22±0.02 |
| Ymax | 98.31 | 99.58 | 94.7±0.51 |
| $R^2$ | 0.9983 | 0.9916 | 0.9947 |
| Sy.x (a) | 1.811 (24) | 3.605 (57) | 2.76 (40) |

The standard deviation of the residuals (Sy.x) = $\sqrt{\frac{\Sigma(residuals^2)}{a-K}}$ where "a" is the number of data points, "K" number of parameters fit by regression and residual is the vertical distance (in Y units) of the point from the fit line.

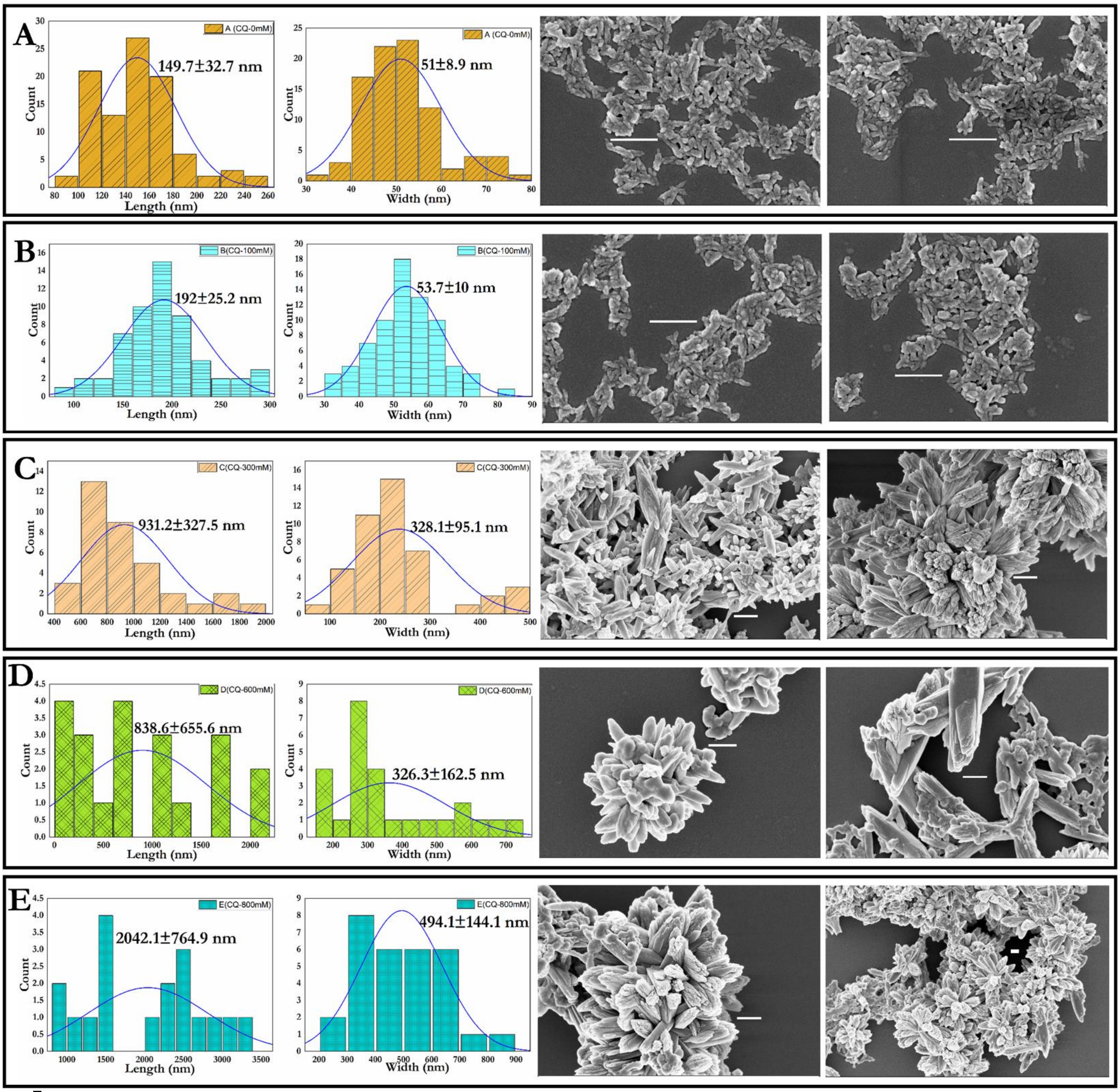


**Figure S5: SEM image analysis:** Particle size distribution histogram determined from the SEM images of β-hematin crystals grown under the same experimental condition, i.e., constant HDP (0.1 µM) and heme concentrations (600 µM) but at different CQ concentrations. A, β-hematin crystals grown in the absence of CQ. Sample B, C, D, and E are the β-hematin crystals grown in the presence of 100 µM, 300 µM, 600 µM, and 800 µM CQ, respectively. The electron micrograph is at different magnifications; therefore, the reference might be of varying size, but they correspond to a common length of 0.5 µm.

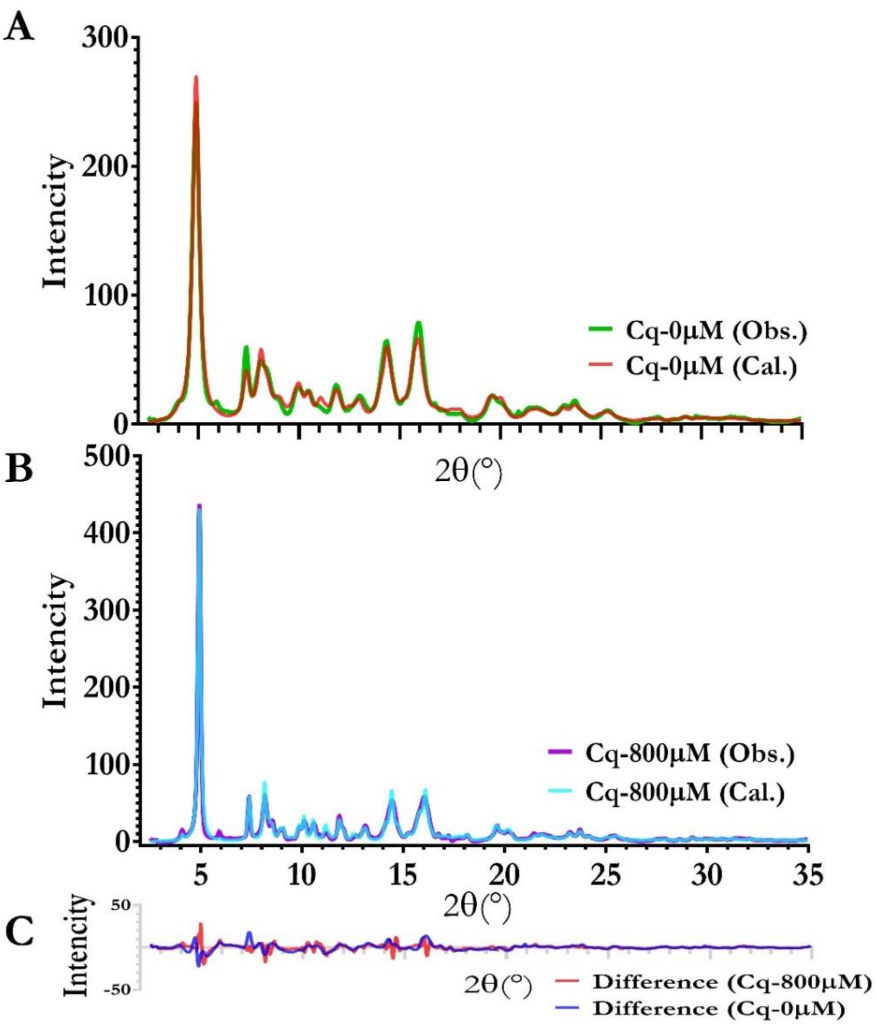


**Figure S6:** Comparison of β-hematin structure from powder XRD: A, B, The color-coded observed (Obs.) and calculated (Cal.) XRD pattern of β-hematin samples. C, color-coded difference curve between observed and calculated XRD pattern intensities. The X-axis of all three graphs (A, B and C) is on the same scale. Unit cell and refinement statics is in table S6

**Table S6:** Unit Cell Dimensions and R Factors for Hemozoin (β-hematin)

| **Parameters** | **Control (Cq-0 µM)** | **Test (Cq-800 µM)** |
|---|---|---|
| Space group | P1 | P1 |
| a (Å) | 12.279 (9) | 12.187 (5) |
| B (Å) | 14.678 (6) | 14.605 (3) |
| C (Å) | 8.031 (4) | 8.038 (2) |
| α (◦) | 90.42 (3) | 90.37 (1) |
| β (◦) | 96.44 (4) | 96.87 (2) |
| γ (◦) | 97.70 (4) | 98.01 (2) |
| Cell Volume | 1425.1 (6) | 1406.35 (3) |
| Crystallite size (nm) | 16.6±1 | 52.5±5 |
| Rwp (%) | 3.70 | 3.33 |

*Figures in parentheses indicate approximate error in the final quoted digit in each quantity
